# Put your money where your mouth is: Surveillance of antibiotic resistance within the commensal *Neisseria*

**DOI:** 10.64898/2026.02.14.705939

**Authors:** Molly R. Regan, Caroline J. McDevitt, Leah R. Robinson, Souwaibat Issifou, Crista B. Wadsworth

## Abstract

Commensal *Neisseria* species are major reservoirs of adaptive genetic variation, including antimicrobial resistance, for their pathogenic relatives, yet they remain poorly characterized. This gap limits our ability to anticipate resistance mechanisms that may ultimately emerge *Neisseria gonorrhoeae* and *N. meningitidis*. Here, we analyzed 166 novel commensal *Neisseria* isolates collected from 31 study participants and measured minimum inhibitory concentrations (MICs) for seven antimicrobials: azithromycin, cefixime, ceftriaxone, ciprofloxacin, doxycycline, and gentamicin. Resistance, defined using the Clinical and Laboratory Standards Institute (CLSI) guidelines, was highly prevalent for azithromycin (76%) and doxycycline (52%), while no resistance to gentamicin was observed. High-level doxycycline resistance was always associated with inheritance of *tetM*. Reduced susceptibility to azithromycin was linked to an MtrD K823E substitution, and reduced susceptibility to ciprofloxacin was associated with GyrA T91I (*N. subflava*) or S91V (*N. mucosa*). The PenA 312M mutation was associated with significantly elevated ceftriaxone and cefixime MICs. Across all antimicrobials, MICs varied widely, indicating the presence of additional modulating mutations. Finally, the genetic determinants underlying low-level doxycycline resistance and reduced penicillin susceptibility remain unresolved. Overall, here we continue to build on the foundation of surveillance efforts in the commensal *Neisseria*, and continue to flesh out what is known and unknown about this early warning system – or canary in the coal mine – for emerging resistance and clinically consequential evolution in pathogenic *Neisseria*.

## Introduction

The rise of antibiotic resistance in *Neisseria gonorrhoeae* represents a critical and escalating threat to global public health. This pathogen, the causative agent of gonorrhea, has demonstrated a remarkable ability to develop resistance to nearly every class of antibiotics used against it, including sulfonamides, penicillin, macrolides, tetracyclines, fluoroquinolones, and, more recently, extended-spectrum cephalosporins^1–3^. As evolving resistance continues to render first-line treatments increasingly ineffective, the risk of untreatable gonococcal infections looms larger, underscoring the urgent need for novel therapeutic strategies and more comprehensive surveillance systems. A key aspect of addressing this crisis lies in understanding the molecular and ecological mechanisms through which *N. gonorrhoeae* acquires antimicrobial resistance (AMR). While clinical surveillance of AMR has historically focused on pathogenic *Neisseria*, a growing body of evidence points to commensal *Neisseria*, harmless colonizers of the oral cavity, as potential and until recently largely overlooked reservoirs of resistance genes^4,5^.

Commensal *Neisseria* species colonize the human naso- and oropharynx. These species are universally carried by healthy human adults and make up the most abundant genera within *Proteobacteria* in oral and pharyngeal samples (approximately 10% of operational taxonomic units)^6,7^. Though species definitions across the *Neisseria* are in flux, the most commonly described human-associated species include those in seven distinct clusters (1) *Neisseria gonorrhoeae*; (2) *N. meningitidis*; (3) *N. lactamica*; (4) the *N. polysaccharea* cluster, which has been proposed to contain multiple species; (5) the *N. subflava* group, which also contains *N. flava* and *N. flavescens*; (6) the *N. mucosa, N.sicca, and N.macacae* group, with a physiologically and genotypically distinct variant *N. mucosa var. heidelbergensis*, and (7) *N. elongata* (atypical rod)^8–11^. More recently, novel species have been proposed, including: *N. benediciae, N. blantyrii, N. bergeri, N. basseii, N. maigaei, N. uirgultaei*, and *N. viridiae*; based on genomic data^8^. However, as noted by Miari et al. (2024)^9^, there are 44 described *Neisseria* species (both human and animal-associated) according to the List of Prokaryotic names with Standing in Nomenclature, and 47 named by the National Center for Biotechnology Information (both databases accessed December 2025); speaking to the complexity and diversity of the genus both phenotypically and genotypically. Interestingly, *Neisseria* species have shown some evidence of localization to distinct ecological niches within the mouth, with *N. meningitidis* and *N. gonorrhoeae* localized to the throat, *N. subflava*, favoring the tongue dorsum, *N. cinerea* preferring the keratinized gingiva and buccal mucosa, and other *Neisseria* mostly localized to the dental plaque^12,13^. However, despite this niche specialization and other barriers to gene exchange (e.g., DNA-uptake sequence divergence^14^, and variation in DNA methylation^15^), DNA donation across species’ boundaries is rampant, and a major factor influencing evolution within the genus^4,5,16,17^.

Closely situated niches in the naso- and oropharynx allows for horizontal gene transfer (HGT) across members of the genus. Genomic DNA flows relatively freely between *Neisseria* species as they constitutively express their competence systems, and remain competent for transformation through all stages of growth. Type IV pili (T4P) promote transformation, and are formed by pilin subunits (PilE), which pass through the outer membrane through the PilQ pore complex^18,19^. Surface-exposed ComP proteins recognize and bind *Neisseria*-specific DNA uptake sequences (DUSs) on extracellular DNA^20^. When the pilus retracts, it draws ComP-bound DNA into the cell. Once in the periplasm, single-stranded DNA bound by ComE is imported through the inner membrane and into the cytoplasm through a pore formed by ComA^21,22^. Homologous recombination integrates DNA into the genome via RecA-mediated pathways (RecBCD and RecF)^22,23^. Additionally, the plasmid pConj, which has a high carriage rate across the *Neisseria*, contains all of the necessary machinery for conjugation (i.e. the tra-like genes encoding T4P and T4SS^24–26^), and thus facilitates the spread of both this plasmid and others (i.e., the beta-lactamase containing plasmid p*bla* and the cryptic plasmid pCryp^27,28^) across the *Neisseria*. Through these mechanisms, commensal *Neisseria* have donated alleles encoding resistance to critical antibiotics, including: azithromycin, ciprofloxacin, penicillin, tetracycline, and extended-spectrum cephalosporins^4^. Indeed, mosaic alleles in pathogenic species, including efflux pumps and promoter regions (*mtr*)^29,30^, penicillin-binding protein 2 alleles (*penA*)^31–34^, and *gyrA* mutations^35^, have all been traced back to commensal origins. In addition, commensal-to-pathogen transfer of pConj and *pbla* plasmids, known resistance determinants, has been documented^28^.

Despite their central role in the spread of antimicrobial resistance, commensal *Neisseria* species remain significantly under-sampled in surveillance efforts. For example, of the 63,311 *Neisseria* genomes deposited to NCBI as of December 2025, 48,529 are *N. gonorrhoeae,* and 13,654 are *N. meningitidis*. Thus, only 2% of currently available genomes represent commensals while 98% are of pathogenic sequences. Furthermore, existing studies are often limited in size, geographic diversity, and phenotypic characterization, leaving major gaps in our understanding of the *Neisseria* resistome and its potential to contribute to treatment failures in pathogenic species. Large-scale efforts to collect, identify, and phenotype commensal *Neisseria* isolates are therefore essential to inform surveillance strategies, guide therapeutic development, and preempt resistance outbreaks. Multiple papers have called for expanded *Neisseria* surveillance programs (i.e., such as GISP from the CDC, and the global Gonococcal Antimicrobial Surveillance Program (GASP) and its European version (Euro-GASP) from the WHO) for commensals including: Kenyon and colleagues^36^; Goytia and Wadsworth^5^; and Wadsworth, Goyita, and Shafer^4^; however, currently there are no national or international efforts to do so.

Here, we add to a growing number of studies collecting data on commensal resistance carriage (e.g., ^9,37–42^)In brief, we collected oral *Neisseria* isolates from human participants using non-invasive sampling, characterize them taxonomically using high-throughput sequencing, and profile their susceptibility to a panel of clinically relevant antibiotics. Our findings aim to expand the current understanding of commensal resistance carriage and to contribute valuable data toward surveillance efforts designed to predict and prevent the emergence of resistance in pathogenic *Neisseria* populations.

## Methods

### Collection of novel Neisseria isolates from study participants

The study was approved by the Institutional Review Board of RIT (protocol number: 02030524, date of approval: 30 May 2024), and all experiments were performed in compliance with relevant guidelines and regulations. Study participants (≥□18 years old) were recruited from students, staff, and faculty at the Rochester Institute of Technology’s (RIT) main campus (USA: Rochester, NY) from the summer of 2024 to the fall of 2025. There were no exclusion criteria other than age. Participants were provided written informed consent prior to participation in the study. After consent, participants were asked to complete a voluntary demographic questionnaire.

For bacterial collection, participants agitated the surfaces of their mouths (i.e., teeth, the roof of their mouth, and the insides of their cheeks) for 30 seconds with their tongue, after which they provided a sample of roughly 1 mL of saliva into a sterile tube. Samples were inoculated onto LBVT.SNR — a media developed for the isolation of *Neisseria* commensals^43^ — and grown for 48 h at 30°C. For each participant, up to ten colonies were picked and re-plated on GCB agar plates supplemented with 1% Kellogg’s solution (hereafter GCB-K plates), then incubated in the same conditions. Once sufficient growth of each colony was achieved, samples were stocked in TSB with 20% glycerol and stored at -80℃. The ability of LBVT.SNR to support the growth of commensal *Neisseria* was assessed using known species from the Centers for Disease Control and Prevention (CDC) and Food and Drug Association’s (FDA) Antibiotic Resistance (AR) Isolate Bank. Strains were struck onto LBVT.SNR plates and grown for 24 hours at 30°C, the emergence of colonies or a lawn was coded as evidence of successful growth.

### Antimicrobial susceptibility testing

Frozen stocks of oral *Neisseria* isolates were revived by streaking onto GCB-K agar plates and incubating at 30°C for 24 hours. Cells from overnight plates were suspended in TSB to a 0.5 McFarland standard and inoculated onto a GCB-K plate with an Etest strip. Following 18–24 h of incubation at 30°C, the MICs of each replicate was recorded. We defined resistance using the Clinical Laboratory Standards Institute (CLSI) breakpoints for *N. gonorrhoeae*^44^. Reported MICs are the mode of 2–3 independent tests and were agreed upon by two independent researchers. Antibiotics assessed include: azithromycin, ceftriaxone, ciprofloxacin, doxycycline, penicillin, tetracycline, and gentamicin (Supplemental Table 1).

### Whole genome sequencing

Cell lysis was performed by suspending cell growth from overnight plates in TE buffer (10 mM Tris [pH 8.0], 10 mM EDTA) with 0.5 mg/mL lysozyme and 3 mg/mL proteinase K (Sigma-Aldrich Corp., St. Louis, MO). DNA isolation utilized the PureLink Genomic DNA Mini Kit (Thermo Fisher Corp., Waltham, MA) with RNase A treatment to remove RNA. Isolated DNA was prepared for sequencing using the Nextera XT kit (Illumina Corp., San Diego, CA), and uniquely dual-indexed and pooled. The final pools were subsequently sequenced on the Illumina Novaseq or NextSeq platforms at the Rochester Institute of Technology Genomics Core using V3 600 cycle cartridges (2□×□300 bp) or 1000/2000 P2 XLEAP-SBS Reagent Kit (2 x 300 bp) cartridges respectively. Read libraries were deposited to the NCBI’s Short Read Archive with accessions for each library reported in Supplemental Table 1.

### Bioinformatics

Quality of each paired-end read library was assessed using FastQC v0.11.9^45^. Adapter sequences and poor-quality sequences were trimmed from read libraries using Trimmomatic v0.39, using a phred quality score□<□15 over a 4 bp sliding window as a cutoff^46^. Reads□<□36 bp long, or those missing a mate, were also removed from subsequent analysis. Read libraries were assembled using SPAdes v.3.14^47^. Species identity for read libraries were defined using kraken2^48^, and subsequently confirmed using PubMLST’s *Neisseria* Typing tool (https://pubmlst.org/bigsdb?db=pubmlst_neisseria_seqdef&page=sequenceQuery)^49^ (Supplemental Table 1). Phylogenies were constructed using MLST loci (*abcZ*, *adk*, *aroE*, *fumC*, *gdh*, *pdhC*, and *pgm*) for all collected isolates in addition to 57 reference genomes (Supplemental Table 2). In brief, loci were derived from each genome using blastn^50^. Sequences were then concatenated and the resultant multi-locus fasta file was aligned using MAFFT v.7.0^51^. A maximum likelihood tree was calculated using RAxML v.8.0^52^ and iTOL^53^ was used to visualize and annotate the resultant trees. Blastn^50^ was also used to search for known resistance-associated mutations^1,4^. The gonococcal reference (NC002946) was used to identify orthologous sequence positions across taxa, which was visualized in Geneious Prime^54^

For commensal strains with evidence of polyphyletic species clustering, one isolate was chosen as a representative reference (*N. subflava*: P0006S004, P0009S001, P0014S002, P0019S002, P0033S006, P0036S002, P0037S002; *N. mucosa*: P0005S001, P0009S002) and reads from all isolates within a species cluster collected from the same sample participant were aligned back to the reference using bowtie2 v.2.2.4 using the “end-to-end” and “very-sensitive” options^55^. Pilon v.1.16^56^ was subsequently used to call variants from the reference sequence. All statistical analyses were conducted in R^57^.

## Results

### Participant demographics and minimum inhibitory concentration testing

Of the 31 study participants, 11 were female, 9 were male, and 11 participants did not report. Participant ages ranged from 18 to 36 (mean=21.6). 100% of study participants carried at least one commensal *Neisseria* isolate. Among the commensal isolates tested, resistance was the most prevalent to the antibiotic azithromycin, with 76% of isolates classified as resistant based on CLSI breakpoint guidelines for *Neisseria gonorrhoeae* (MICs ranged from 0.38 to 256 µg/ml, Table 1). Resistance to doxycycline was the next most prevalent, with 52% of isolates found to be resistant (MICs: 0.023-256 µg/ml, Table 1). For the β-lactams, we found 9% resistant to penicillin (MICs: 0.0125-4 µg/ml), 7% resistant to cefixime (MICs: 0.016-4 µg/ml), and 3.8% resistant to ceftriaxone (MICs: 0.008-1.5 µg/ml). We found only 2.5% of isolates resistant to ciprofloxacin (MICs: 0.002-2 µg/ml) and no isolates with resistance to gentamicin (MICs: 0.75-8 µg/ml).

**Table 1.** Summary of antimicrobial resistance within the commensal *Neisseria* isolates collected for this study.

| Antibiotic | N | MIC Min<br>(µg/ml) | MIC Max<br>(µg/ml) | MIC Mean<br>(µg/ml) | MIC Median<br>(µg/ml) | SD | No.<br>Resistant | Resistant<br>(%) |
| --- | --- | --- | --- | --- | --- | --- | --- | --- |
| Azithromycin | 157 | 0.38 | 256 | 9.24 | 3 | 31.56 | 119 | 76 |
| Cefixime | 151 | 0.016 | 4 | 0.14 | 0.064 | 0.38 | 11 | 7 |
| Doxycycline | 157 | 0.023 | 256 | 14.75 | 1 | 39.6 | 81 | 52 |
| Penicillin | 154 | 0.125 | 4 | 0.99 | 1 | 0.63 | 14 | 9 |
| Ceftriaxone | 157 | 0.008 | 1.5 | 0.08 | 0.047 | 0.14 | 6 | 3.8 |
| Ciprofloxacin | 157 | 0.002 | 2 | 0.12 | 0.012 | 0.3 | 4 | 2.5 |
| Gentamicin | 155 | 0.75 | 8 | 2.2 | 2 | 0.88 | 0 | 0 |

AZI ≥2, PEN ≥ 2, CRO > 0.25, CFX > 0.25, CIP ≥ 1, DOX ≥ 1\*, GEN ≥ 16\*
\*Doxycycline and gentamicin have no established clinical breakpoints for *Neisseria*

### Genomics data

In total, 166 samples were sequenced. The mean genome length was 2.44 Mb, with a minimum of 2.13 Mb and a maximum of 3.00 Mb. Kraken2-based species determination identified *N. subflava* to be the predominant species present amongst the study participants (121/165, 73%, Figure 1A). The second most commonly carried species was *N. mucosa* (40/165, 24%), followed by *N. elongata* (2/165, 1%) and unclassified *Neisseria* species (2/165, 1%). After excluding the two participants with only a single isolate collected (P0012 and P0023), 14 participants carried a single *Neisseria* species (*N. subflava,* 48.2%), 14 carried two species (*N. subflava* and *N. mucosa*, 48.2%), and 1 participant carried 3 (P0008, the only parcicpant to carry *N. elongata*). (Figure 1B). Looking at MIC distributions across species, we note that *N. subflava* isolates had the highest proportion of resistant isolates for all antimicrobials: Azithromycin 74% (89/119), ceftriaxone 100% (6/6), cefixime 100% (11/11), ciprofloxacin 75% (3/4), doxycycline 74% (60/81), and penicillin 50% (7/14) (Figure 2).

**Figure 1.**
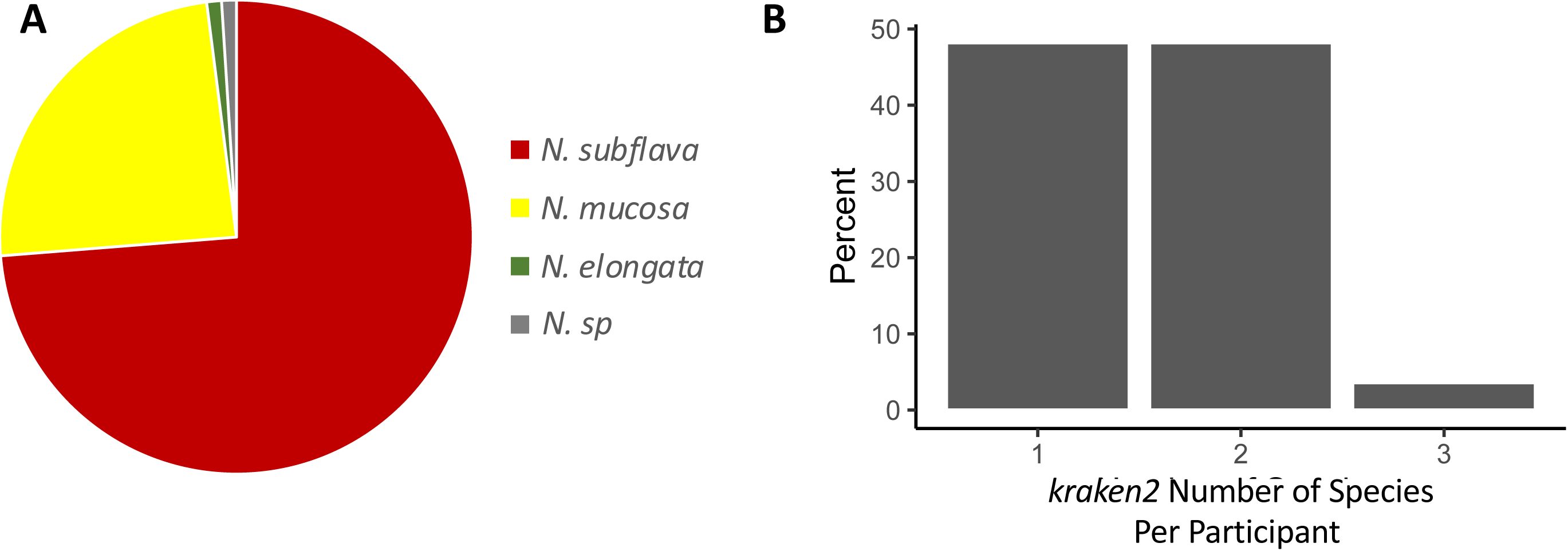
Neisserial species diversity for 166 isolates collected from Rochester, NY between 2024-2025. (A) Kraken2 identified *N. subflava* to be the main species present amongst the study participants (121/165, 73%), followed by *N. mucosa* (40/165, 24%), *N. elongata* (2/165, 1%), and unclassified *Neisseria* species (2/165, 1%). (B) Excluding participants with only a single isolate collected (P0012 and P0023), fourteen participants carried a single *Neisseria* species (*N. subflava,* 45%), while the remainder carried more than one. No participant carried more than three species.

**Figure 2.**
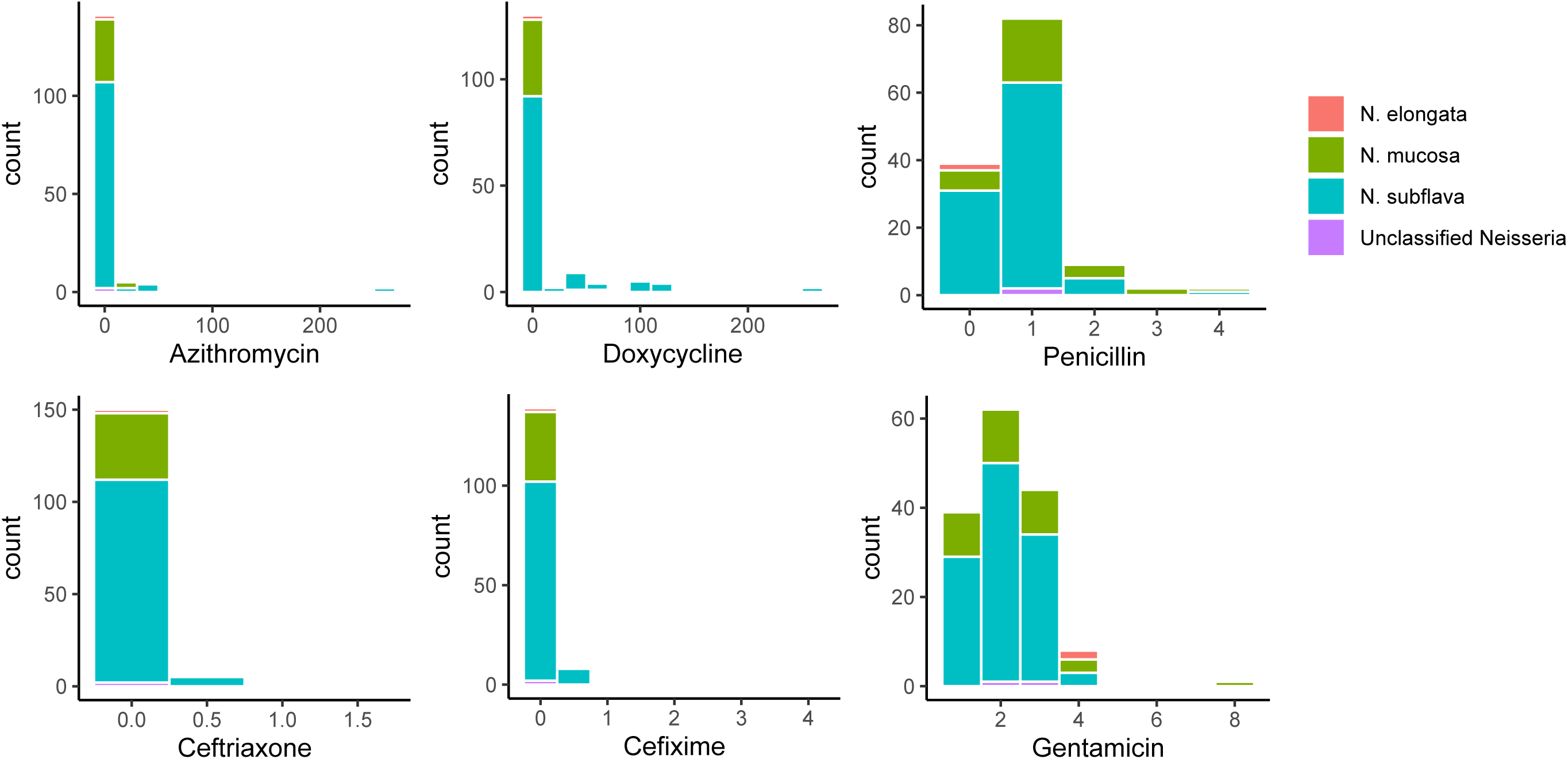
Minimum inhibitory concentration (MIC) distributions across *Neisseria* species for (A) azithromycin, (B) doxycycline, (C) penicillin, (D) ceftriaxone, (E) cefixime, and (F) gentamicin. *N. subflava* made up the highest percentage of resistant isolates for all antimicrobials: Azithromycin 74% (89/119), ceftriaxone 100% (6/6), cefixime 100% (11/11), ciprofloxacin 75% (3/4), doxycycline 74% (60/81), and penicillin 50% (7/14).

Phylogenetic clustering confirmed kraken2 species’ identifications, and clarified the two unclassified *Neisseria* belonged to the *N. subflava* clade (Figure 3). This analysis also confirmed that we did not collect any *Neisseria* from *N. cinerea, N. polysaccharea*, newly nominated *Neisseria* species^8^, or the pathogenic *Neisseria* clade (*N. meningitidis*, *N. gonorrhoea*, and closely related non-pathogenic *N. lactamica*) (Figure 3). As a note, *Neisseria* taxonomy changes substantially depending on the group, analysis, and dataset. As per recent reports, we merged the *N. subflava* and *N. flavecens* species into a single group/cluster^9,58^; and merged the *N. mucosa, N. macacae*, and *N. sicca* species into a single cluster^9,58^. Reference genomes for these species were polyphyletic within groups supporting this merging strategy. For the *N. subflava* group, 31 participants carried this species, with six participants showing evidence of carrying at least two phylogenetically distinct strains (Figure 4A). For the *N. mucosa* group, 15 participants carried this species, with two participants carrying at least two phylogenetically distinct strains (Figure 4B). The two *N. elongata* strains were collected from the same participant, and appear to be closely related; though with limited strains, overall diversity between these two isolates is not clear.

**Figure 3.**
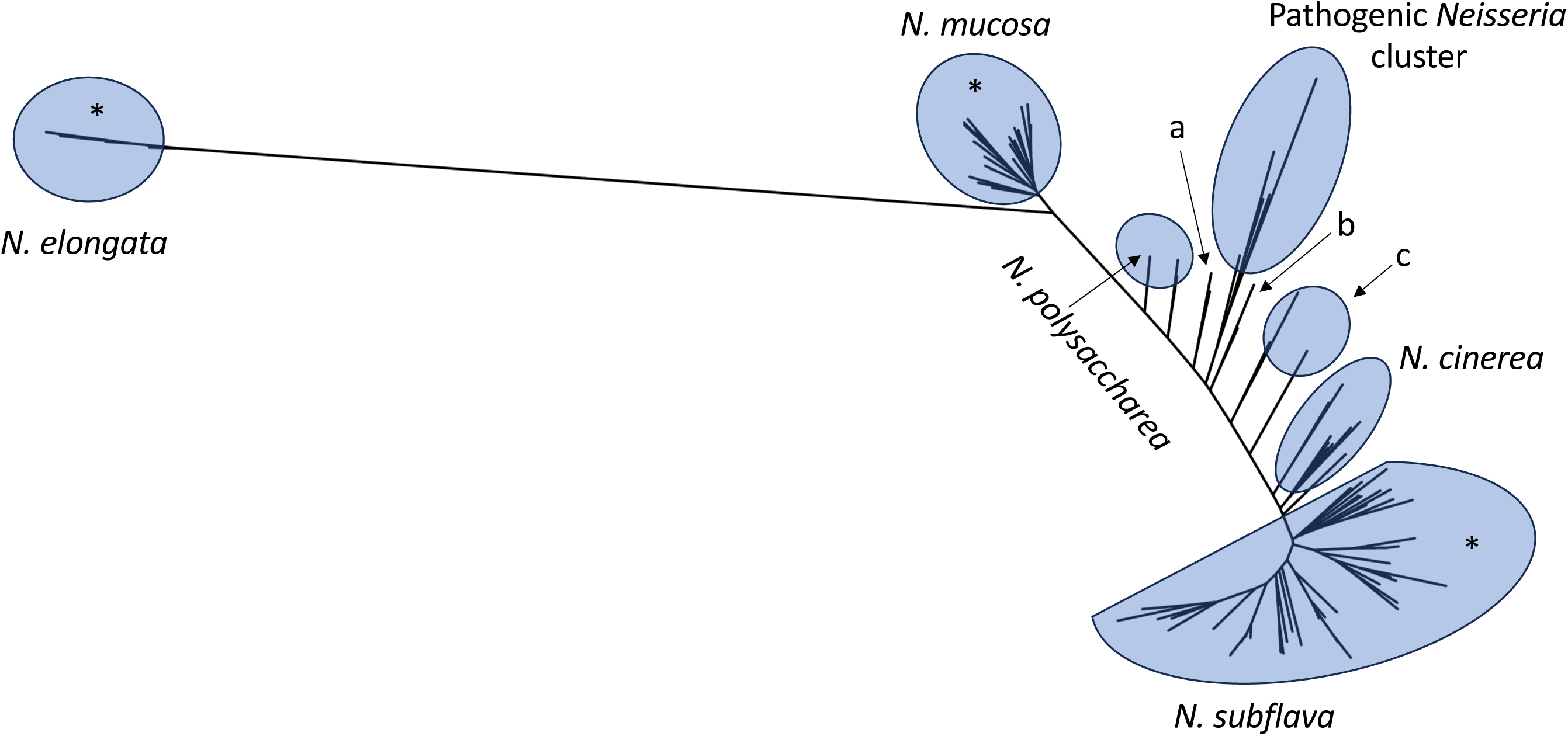
MLST gene-based maximum likelihood phylogeny for the 166 isolates collected within the study and 57 *Neisseria* reference isolates. Isolates recovered from participants on LBVT.SNR media included those clustering within the *N. elongata, N. mucosa*, and *N. subflava* species groups (indicated by *). We did not collect any isolates from the *N. cinerea* cluster, polyphyletic *N. polysacchaera* group, pathogenic *Neisseria* cluster (including *N. gonorrhoea*, *N. lactamica*, N. maigaei, *N. meningitidis*, and *N. uirgultaei*), (a) *N. viridiae* group, (b) *N. benedictiae* and *N. blantyrii* group, or (c) *N. basseii* and *N. bergeri* polyphyletic cluster.

**Figure 4.**
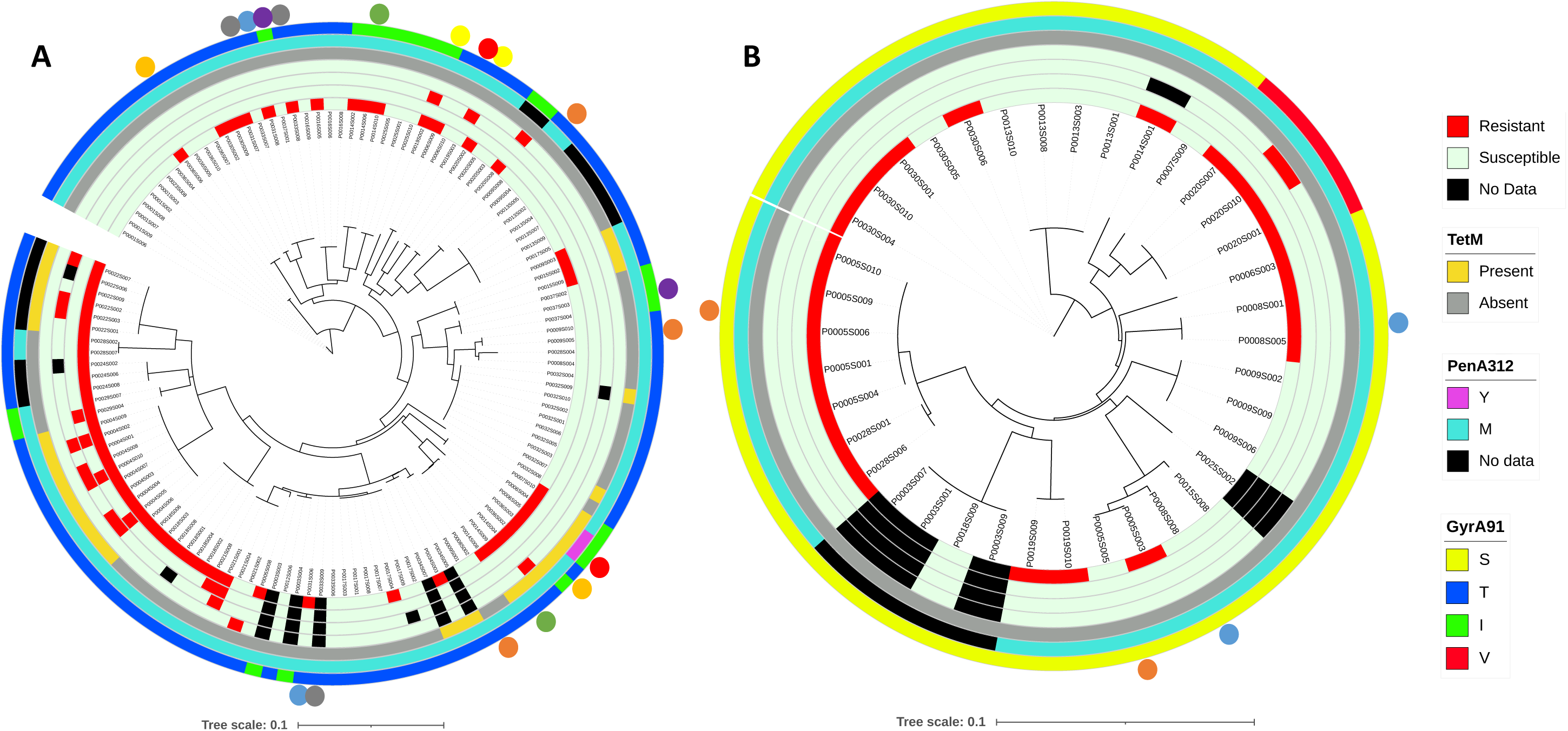
MLST maximum likelihood phylogenies by species for (a) *Neisseria subflava* and (b) *N. mucosa.* Annotation rings from the center indicate resistance (red) or susceptibility (green) for doxycycline, ceftriaxone, cefixime, and ciprofloxacin. The fifth annotation ring indicated presence (yellow) or absence (grey) of the *tetM* gene, followed by annotation rings indicating amino acids present at PenA312 and GyrA91. Colored circles on the outermost annotation rings indicate strains within participants that appear to be genetically distinct (i.e., clustering in multiple regions of the phylogeny).

Blast was used to search for known resistance-associated mutations^4^, using gonococcal sequence positions as a reference (NC002946, Supplementary Table 1). High-level tetracycline and doxycycline resistance was always imparted by inheritance of the pConj plasmid containing the ribosomal protection protein TetM, which we observe in 32 samples. Isolates with *tetM* had significantly higher doxycycline MICs than those isolates that did not have evidence of carrying the gene (T-Test: P <0.00001; Figure 5A). MtrR A39T and G45D mutations can also contribute^59^, however were not observed. Finally, elevated MICs can also be imparted by RpsJ V57M^60^, however, samples were monomorphic for wild-type alleles. Ciprofloxacin resistance has been associated with the GyrA substitutions: S91F and D95A/N/G^61,62^. There was variability in our dataset at position 91 with isolates harboring either S (n=37), V (n=3), T (n=105), or I (n=21) substitutions; yet all samples were wild-type at position 95. Isolates harboring a V or an I substitution at position 91 had significantly higher MICs than those with an S or a T (Tukey’s HSD: P<0.00001 in all cases; Figure 5B). Resistance-associated MtrR A39T, G45D and Y105H^63^ were not found. Finally, the resistance-associated GyrB mutation D429N was not found which imparts cross-resistance to zoliflodacin and gepotidacin, as well as other zoliflodacin-resistance associated mutations D429N and S467N^64,65^. The *23S rRNA* C2611T and A2059G^66^, MtrR A39T and G45D^59^, MtrD K823E^29,30^, RplD G68D/C and G70D^67^, RplV^68^ and RpmH^69^ tandem duplications have all been associated with elevated MICs to azithromycin in prior studies. Here, we found no evidence of variation at *23S* 2611 or 2059, MtrR at amino acid position 39, and RplD 68 or 70; or RplV or RpmH tandem duplications. Interestingly, all isolates had a G45A substitution in MtrR, however this was not associated with elevated MICs. At MtrD only two isolates (both *N.elongata*) had the wild-type K amino acid at position 823, however four isolates had MtrD K823S, and 160 had K823E. Though MICs for isolates with 823 S or E appeared elevated, they were not significantly different from wild-type (Figure 5C). Gentamicin resistance has been associated with mutations in FusA A563V, G564D, V651F^70^; however, all were homogenous for the wild-type allele in our dataset.

**Figure 5.**
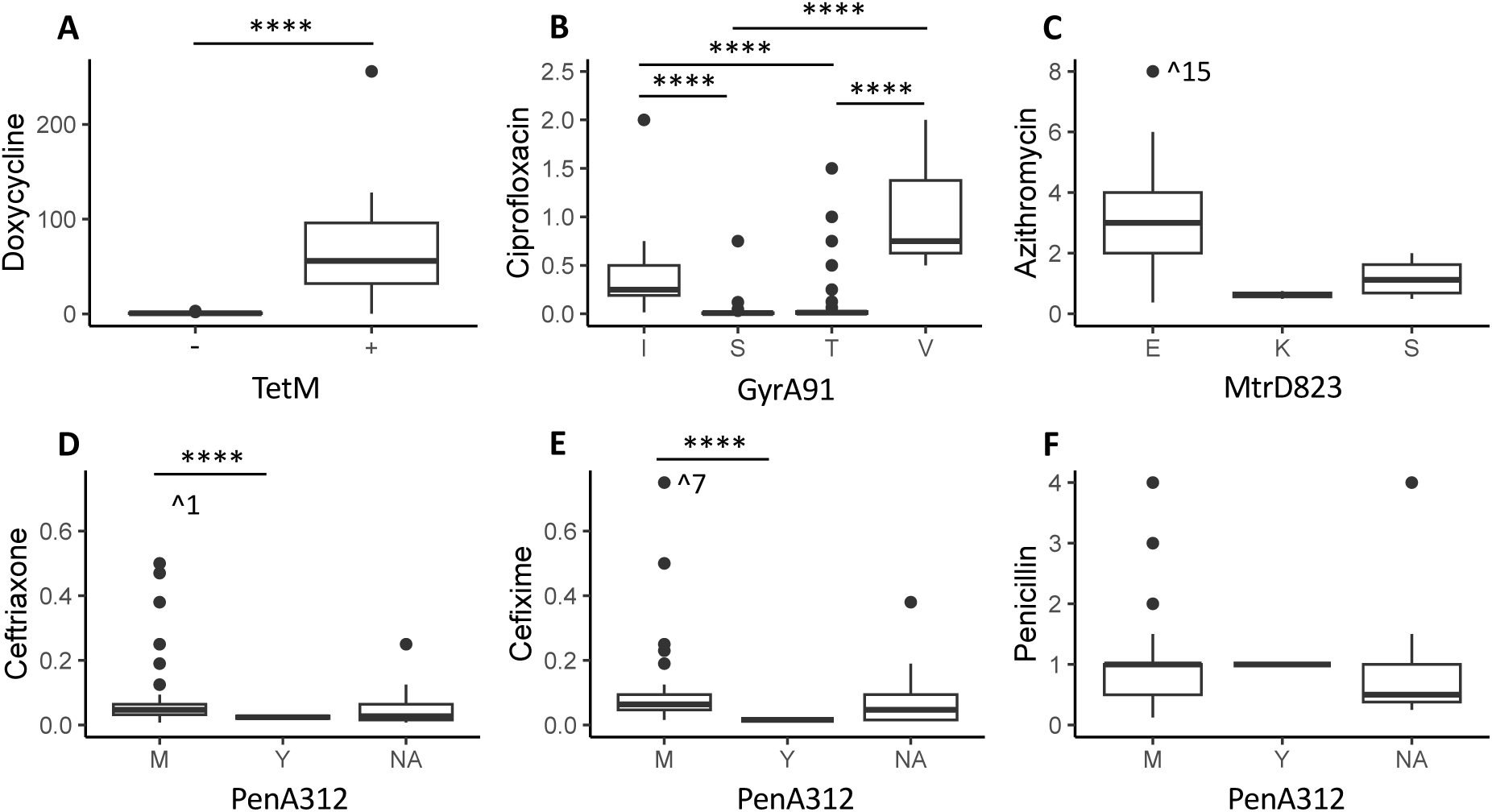
Associations between resistance mutations and MICs for isolates collected within the study. (A) High doxycycline MICs were significantly correlated with inheritance of the *tetM* gene. (B) Isoleucine and serine substitutions at GyrA position 91 resulted in significantly higher ciprofloxacin MICs. (C) Isolates inheriting MtrD823E had higher azithromycin MICs, however this was not a significant difference. (D-F) PenA312M was associated with higher ceftriaxone and cefixime MICs, however not higher penicillin MICs.

For cephalosporin resistance, there are many amino acids that have been suggested to play a role in elevated MICs^71,72^; we searched for eight: A311V, I312M, T316P, T483S, A501V, F504L, N512Y, and G545S. Of all of these positions, the only variation was found at position 312, with 141 isolates having an M substitution and 2 with a Y. For isolates for which we had sequence data, inheritance of the M mutation was associated with significantly higher ceftriaxone MICs (T-test: P<0.00001; Figure 5D) and cefixime MICs (T-test: P<0.00001; Figure 5E), but not penicillin MICs (T-test: P=0.76; Figure 5F). All isolates for which we had sequence data were fixed for the L substitution at position 504 (n=141); and all other amino acids were wild-type. RpoB R201H and RpoD E98K and Δ92–95 have also been associated with higher ceftriaxone MICs^73^, however these mutations were not found. Penicillin resistance can be caused by either chromosomal or plasmid-based mutations^74^. We did not detect the β-lactamase containing plasmid (p*bla*) in any of the samples reported in this study. Chromosomal mutations contributing to resistance include the aforementioned *penA* and *mtrR* mutations, as well as PonA L421P which was not detected.

Carriage of resistance-associated mutations across species was distinct (Figure 4). The *tetM* gene was only found in *N. subflava* and not *N. mucosa* isolates. Carriage of tetM was associated with resistance in all cases except for two isolates which carried *tetM* but had MICs of 0.25 (P0032S009) and 0.5 µg/ml (P0032S007). MICs for *N. subflava* isolates outside of these two ranged from 1 to 256 µg/ml. Though the 312M substitution was associated with elevated ceftriaxone and cefixime MICs (Figure 4D-4E), resistance above the breakpoints of > 0.25 µg/ml was not consistently associated with inheritance of this mutation in either species (Figure 4). Resistance to both these extended spectrum cephalosporins was mostly found in *N. subflava*, with only one *N. mucosa* isolate (P0020S010) with an MIC above the breakpoint value (Figure 4). Finally, GyrA alleles were distinct across species, with *N. subflava* carrying GyrA 91 T and I, and *N. mucosa* carrying S and V. Inheritance of I or V was associated with significantly higher MICs (Figure 4B). *N. elongata* isolates were susceptible to all drugs investigated and only carried mutations associated with lower MIC values (Supplemental Table 1).

## Discussion

Commensal *Neisseria* species remain profoundly under sampled and under characterized relative to their pathogenic counterparts, despite mounting evidence that they play a critical role in the emergence and dissemination of antimicrobial resistance across the genus^4,5^. These organisms serve as a substantial and dynamic reservoir of resistance determinants, yet they are rarely included in routine surveillance or large-scale genomic studies^5^. In this work, we directly address this gap by expanding the catalog of characterized *Neisseria* commensals. We collected and curated a set of novel commensal isolates and systematically surveyed them for antimicrobial resistance phenotypes and their associated genetic determinants. By pairing resistance profiling with genomic analysis, we aim to define both the breadth of resistance present in these species and the specific alleles that may be mobilized through horizontal gene transfer.

Reports of human colonization by commensals vary, ranging from 10.2-100%^9,42,75–77^. However, as previously discussed by Miari et al.^9^, this is likely due to the sampling strategy and/or media used. For example, studies highlighting a lower colonization rate used pharyngeal swabbing and modified Thayer Martin (TM) agar plates^75^, which both will select against commensals due to ecological niche specificity of these species (i.e., commensals are less frequently found in the pharyngeal niche compared to other oral sites^12,13^) and the design of TM plates for isolation of pathogenic *Neisseira*. We add to a growing number of studies which support a 100% carriage rate for commensals^76^. Species diversity however was low, with most isolates being *N. subflava* (123/165, 75%), followed by *N. mucosa* (40/165, 24%), then *N. elongata* (2/165, 1%); which we hypothesized was most likely due to exclusive usage of LBVT.SNR media and growth conditions at 30℃ in a non-CO2 incubator, as at least the pathogenic *Neisseria* require nutrient rich media, and prefer higher temperatures and a 5% CO2 atmosphere^78^. Commensals more closely related to the pathogenic *Neisseria* (Figure 3), also failed to grow in the conditions used within this study. However, after testing the growth of known commensal strains on LBVT.SNR at 30℃ we observed evidence the culture conditions and media supported growth of these other species (Supplementary Table 3). Therefore, limited species diversity may be more likely to be due to the spit collection protocol. If *Neisseria* inhabit distinct niches within the oral cavity as some studies suggest^12,13^ it may be difficult to sample some species and sites (e.g., keratinized gingiva and/or gingival sulcus for *N. cinerea*) using a straightforward spit protocol; swabbing of these sites may in fact be a more appropriate in these cases.

Investigation of interspecific diversity within participants suggests that most participants carried predominantly *N. subflava* (48%) or a mix between *N. subflava* and *N. mucosa* (48%). *N. elongata* was rare and only isolated from one participant, P0008, however interestingly that participant also had the highest interspecific diversity, carrying all three isolated species. Although characterized using different methodology (i.e., phenotypic rather than genotypic species identification) other studies also find the majority of people carry between 1-3 commensal *Neisseria* species using a similar collection protocol^9^. Full analysis of species carriage diversity will likely have to use multiple collection protocols, due to the specific growth requirements of diverse *Neisseria* species, and site specificity as discussed above.

Within host strain diversity has been assessed previously in *N. gonorrhoeae*^79,80^ and *N. meningiditis*^81,82^, however these studies often focus on evolution across the course of colonization or infections, or diversity between isolates colonizing different body sites in the case of *N. gonorrhoea*^83,84^. Here, we assess intraspecific variation between isolates collected at the same time from the same location within individuals in *Neisseria* commensals. We find evidence of 8 participants with at least two genetically distinct strains of *N. subflava* (P0006, P0009, P0014, P0019, P0031, P0033, P0036, and P0037) and 2 participants with at least two genetically distinct strains of *N. mucosa* (P0005 and P0009); as indicated by polyphyletic clustering patterns (Figure 4). For these participants, *N. subflava* isolates carried on average 65,430 single nucleotide polymorphisms (SNPs) compared to the participant-specific reference (Figure 6A); and *N. mucosa* strains were separated by on average 28,927 SNPs (Figure 6B). Overall, genomes ranged from 0.0009-3.65% divergent compared to reference sequences; with strains P0005S003 and P0005S005 on the high end of the range, differing by >100,000 SNPs compared to the reference (P0005S001). Generally, for bacteria, an average nucleotide identity (ANI) of ≥95% is used to denote distinct species, so while our data suggests the possibility of distinct strains within-hosts there is not enough divergence in this dataset to suggest lineage splitting. Two possibilities exist for the observation of intraspecific within-host diversity: 1) recent colonization of hosts with novel strains, with previous studies finding that commensals are shared between close contacts^85^, or 2) within-host oral colonization site divergence within a species, which has been demonstrated between species^12,13^, but not within a species at this point.

**Figure 6.**
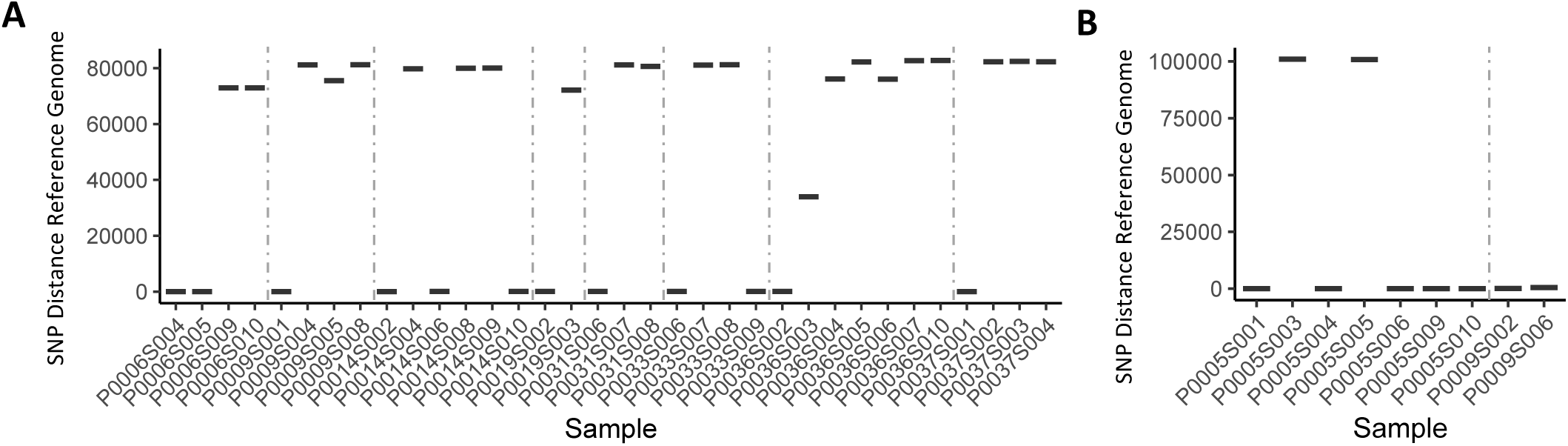
Within-host variation of *Neisseria* commensals indicates intraspecific divergence. (A) For *N. subflava*, 8 participants carried at least 2 distinct strains. These strains carried on average 65,430 single nucleotide polymorphisms (SNPs) compared to the participant-specific reference. (B) For *N. mucosa*, 2 participants carried at least 2 distinct strains separated by on average 28,927 SNPs. Overall, genomes ranged from 0.0009-3.65% divergent compared to reference sequences.

Surveillance of commensals may serve may to ultimately inform on circulating resistance determinants, known or unknown, that may disseminate into pathogenic *Neisseria* populations as has been observed for other resistance markers before^4,5^. In this dataset, resistance was highly prevalent for azithromycin (76%) and doxycycline (52%), and while no resistance to gentamicin was observed, resistance was present to all other drugs investigated. As azithromycin and doxycycline are among the most frequently prescribed oral antibiotics prescribed in the United States^86^, this may suggest high use is selecting for elevated resistance carriage in commensal populations through bystander selection. Indeed, we have previously reported that doxycycline use is directly correlated with doxycycline resistance carriage and elevated doxycycline MICs in *Neisseria*^63^, which we confirm in this expanded dataset (Supplementary Figure 1). High-level doxycycline resistance > 8 µg/ml was always associated with inheritance of the *tetM* gene (Figure 5A), while the mutations imparting low-level resistance are unclear as RpsJ 57 mutations were not present. The prevalence of doxycycline resistance, and of the conjugative plasmid (pConj) harboring *tetM* is of course concerning due to the implementation of doxy-PEP^87,88^. We can only anticipate that doxy-PEP selection will further spread the highly prevalent *tetM* and pConj in commensal populations, which has been predicted in mathematical models for *N. gonorrhoeae*^89^. Furthermore, increased prevalence of pConj may also spread β-lactam resistance via transfer of p*bla*. Ultimately, doxy-PEP will likely increase the available pool of these plasmids and their resistance determinants to the pathogenic *Neisseria*.

Reduced susceptibility to ciprofloxacin in *N. gonorrhea* is frequently associated with GyrA S91F and D95A/N/G substitutions^61,62^. We did not observe any variability at position 95, however there was variability in our dataset at position 91 with isolates harboring either S (n=37), V (n=3), T (n=105), or I (n=21) substitutions. Reduced susceptibility was associated with GyrA T91I (*N. subflava*) or S91V (*N. mucosa*) (Figure 5B). In a previous report investigating alleles in the PubMLST database, we find that 3% of commensals harbor the resistance-associated GyrA 91I substitution^90^, suggesting its availability in natural commensal populations. The presence of additional mutations were also noted: Reduced susceptibility to azithromycin was linked to an MtrD K823E substitution previously reported to contribute to resistance in *N. gonorrhoeae* isolates harboring mosaic *mtr* alleles^29,30^ (Figure 5C), and a PenA 312M mutation was associated with significantly elevated ceftriaxone and cefixime MICs^71,72^ (Figure 5C). Importantly, across all antimicrobials, MICs varied widely, likely indicating the presence of additional modulating mutations; and finally, the genetic determinants underlying low-level doxycycline resistance and reduced penicillin susceptibility remain unresolved.

In the absence of coordinated national or international strategies for commensal surveillance, the responsibility for documenting these species currently rests with individual laboratories. While fragmented, these efforts are essential. Together, they provide an early warning system, effectively a canary in the coal mine^5^, potentially revealing resistance alleles circulating silently in nonpathogenic populations before they are detected in clinically significant pathogens. Ultimately, a more comprehensive understanding of commensal *Neisseria* (i.e., Keyon et al.’s proposed ‘pan-*Neisseria*’ approach^36^) will illuminate the full resistome accessible to the genus as a whole and identify genetic variants that may be poised for rapid transfer into pathogenic species. Such knowledge is crucial for anticipating future resistance trajectories and informing proactive antimicrobial stewardship and surveillance efforts.

## Acknowledgements

The authors would like to acknowledge the generous support provided by the RIT College of Science and the Thomas H. Gosnell School of Life Science for this work. Work reported in this publication was also supported by the National Institute of General Medical Sciences of the National Institutes of Health under Award Number R15AI174182. The content is solely the responsibility of the authors and does not necessarily represent the official views of the National Institutes of Health. The funders had no role in study design, data collection and analysis, decision to publish, or preparation of the manuscript. The authors would also like to thank Girish Kumar at the RIT Genomics Core for providing support and sequencing services.

---

**Supplementary Figure 1. Doxycycline use and correlation with elevated doxycycline MICs.** Use was significantly associated with elevated MIC values.

