## Supplementary figures and images for "Put your money where your mouth is: Surveillance of antibiotic resistance within the commensal *Neisseria*"

### Supplementary Figure 1

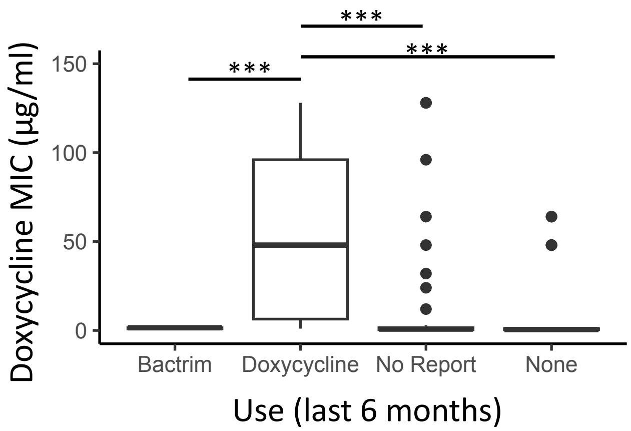
